# Methylmalonic acid: a new target for Hadamard-edited MRS

**DOI:** 10.64898/2026.06.05.730405

**Authors:** Yulu Song, Tao Gong, Zahra Shams, Xiaoya Sun, Christopher W. Davies-Jenkins, Shuyuan Wang, Gizeaddis L. Simegn, Saipavitra Murali-Manohar, Abdelrahman Gad, Georg Oeltzschner, Guangbin Wang, Richard A.E. Edden

## Abstract

**Background:** Methylmalonic acidemia (MMAemia) is a genetic metabolic disorder characterized by an accumulation of methylmalonic acid (MMA) and impaired energy metabolism leading to increased lactate (Lac). The signals of MMA (1.23 ppm) and Lac (1.33 ppm) overlap, making their separation using conventional MRS challenging. An MRS method to differentiate the two metabolites could enhance pathophysiological understanding and improve treatment monitoring - Hadamard-edited MRS has the potential to achieve this.

**Purpose:** To develop a Hadamard-encoded *J*-difference editing approach for independent detection of MMA and Lac at 3T.

**Methods:** A novel Hadamard-encoded editing scheme was implemented and evaluated with density-matrix simulations, phantom and *in vivo* experiments. The new four-step scheme uses frequency-selective editing pulses, applied at 3.2 ppm and 4.1 ppm to modulate the *J*-coupled methyl resonances of MMA and Lac, respectively. Hadamard combinations of the four sub-experiments yield the separate difference-edited spectra for each target metabolite.

**Results:** Simulations and phantom experiments clearly illustrate the separated signals of MMA and Lac. *In vivo* validation experiments show a Lac signal (but no MMA) in a healthy infant, and both Lac and MMA (separated into their respective Hadamard-combination spectra) in a patient with MMAemia.

**Conclusion:** Hadamard-encoded editing at 3T can separate MMA and Lac signals and shows promise for studying altered metabolism in patients with MMAemia.

## 1. Introduction

Methylmalonic acid (MMA) is an intermediate in the metabolic pathway that converts propionyl-coenzyme A (propionyl-CoA) to succinyl-coenzyme A (succinyl-CoA), which then enters the tricarboxylic acid (TCA) cycle for energy production^1,2^. It is normally present at *very* low levels in the brain, but increased levels can arise due to genetic mutations in the propionate pathway, vitamin B_12_ deficiency, and aggressive cancers^3^. MMA is recognized as a significant pathogenic biomarker associated with methylmalonic acidemia (MMAemia), a rare neurometabolic disorder. Accumulation of MMA in MMAemia is caused by a deficiency of methylmalonyl-CoA mutase, which catalyzes the conversion of methylmalonyl-CoA to succinyl-CoA^1^. MMA accumulation causes brain tissue damage, which can manifest as various degrees of intellectual disability^3^ and severe neurological dysfunction. The mechanisms underlying MMA-induced brain damage are multifactorial and include mitochondrial dysfunction characterized by energy failure and oxidative stress^4,5^, hypomyelination and demyelination responses, as well as excitotoxicity resulting from neurotransmitter dysregulation^6^.

Proton magnetic resonance spectroscopy (^1^H-MRS) is a powerful, widely used non-invasive technique measuring brain metabolite levels *in vivo*. It is limited by incomplete separation of metabolite signals, due to limited chemical shift dispersion, broad linewidths in living tissue, and signals split into multiplets by scalar coupling. As a result, signals from low-concentration metabolites are frequently masked by stronger signals, making accurate quantification difficult. Spectral editing techniques, such as MEscher-GArwood Point RESolved Spectroscopy (MEGA-PRESS^7^), enable the selective detection of signals from desired metabolites by selectively manipulating signals of interest and subtracting overlapping signals from more abundant metabolites.

Hadamard Encoding and Reconstruction of MEGA-Edited Spectroscopy (HERMES)^8^ was developed to address one drawback of MEGA-PRESS — that it only edits one metabolite at a time. By using dual-band editing pulses simultaneously applied to distinct editing targets, HERMES can simultaneously and separably detect more than one metabolite with overlapping signals within a single acquisition (e.g., GABA and GSH^9^).

The MR spectrum in MMAemia often shows increased signal around 1.3 ppm that has historically been assigned to lactate (Lac)^10^. The doublets of MMA (1.23 ppm) and Lac (1.33 ppm) overlap at 3T, making their separation using conventional MRS a challenge. *J*-coupling effects in MMA and Lac further complicate the evolution and separation of their signals. The respective coupling partners (adjacent protons within the same molecule) of these two overlapping resonances are well resolved (i.e., MMA at 3.2 ppm and Lac at 4.1 ppm), so editing pulses can be independently applied to each. Therefore, a novel four-step HERMES editing scheme was proposed, enabling independent manipulation of the MMA and Lac spin systems. Applying a Hadamard transformation to the acquired data produces distinct difference-edited spectra for each metabolite. We validate this scheme using density-matrix simulations, phantom measurements, and *in vivo* studies in a healthy control and a patient with MMAemia (3 brain regions each).

## 2. Methods

### 2.1 HERMES editing of MMA and Lac

*J*-difference editing involves the acquisition of two sub-experiments (referred to as ON and OFF), the first of which applies a frequency-selective editing pulse to the target spin system. Subtraction of these sub-experiments yields a difference spectrum that only contains those signals that are affected by the editing pulse. HERMES is a multiplexed implementation of *J*-difference editing that allows simultaneous editing of more than one metabolite. Provided that the editing target signals are sufficiently well resolved, editing pulses can be independently applied to them - in this case, to MMA spins at 3.2 ppm and Lac spins at 4.1 ppm, as shown in Figure 1a. Four sub-experiments (labeled A, B, C and D, respectively) are performed that apply editing pulses to: both Lac and MMA (ON_Lac_, ON_MMA_); MMA only (OFF_Lac_, ON_MMA_); Lac only (ON_Lac_, OFF_MMA_); or neither (OFF_Lac_, OFF_MMA_), in an interleaved fashion. The Lac-edited difference spectrum is calculated by subtracting the two OFF_Lac_ sub-experiments from the sum of the two ON_Lac_ sub-experiments. Similarly, the sum of the two ON_MMA_ sub-experiments minus the sum of the two OFF_MMA_ sub-experiments gives the MMA-edited difference spectrum.

**Figure 1.**
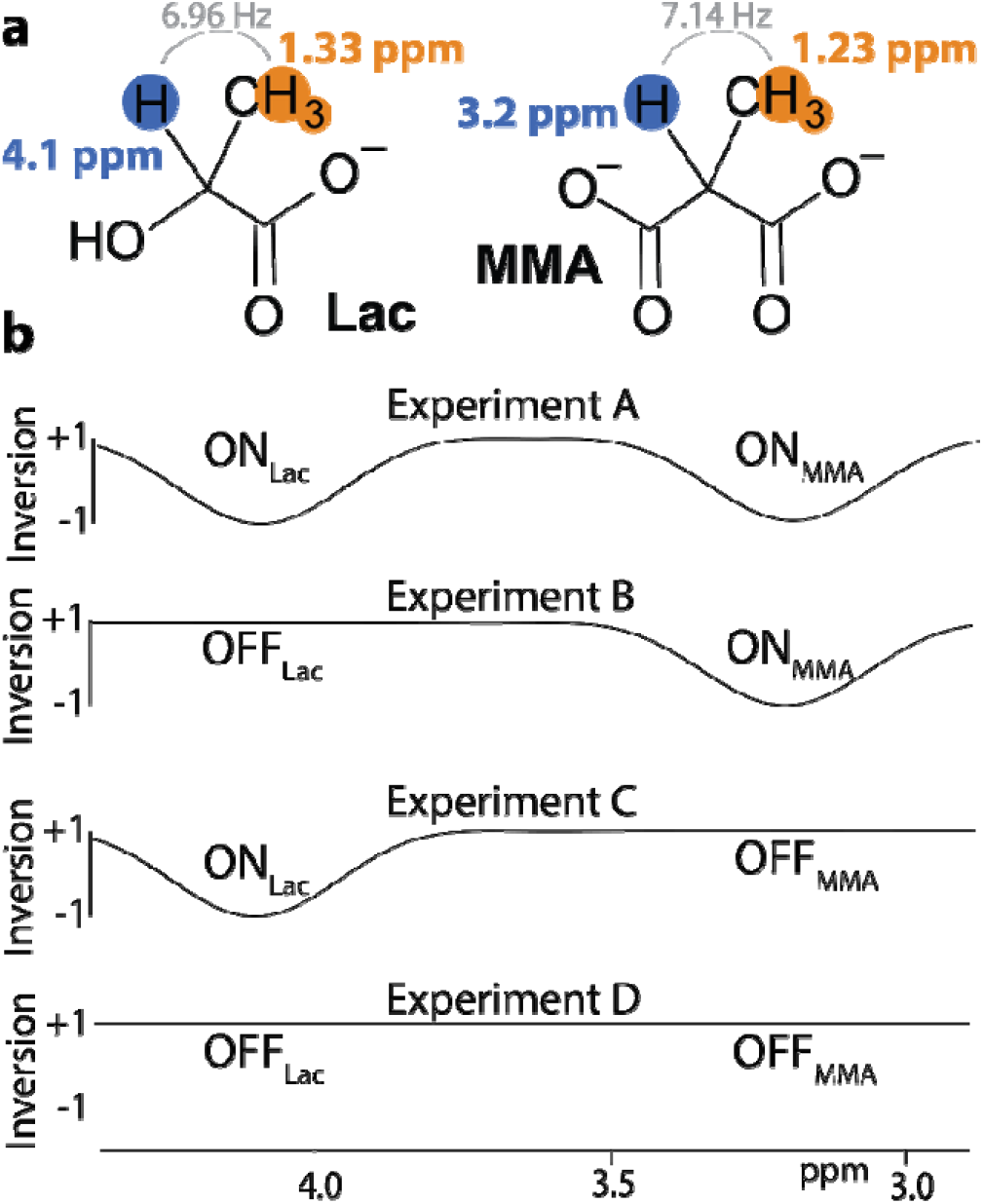
a. Chemical structures of Lac and MMA, including the spin-system parameters of their respective methyl moieties, which are targeted by the HERMES experiment. b. HERMES editing scheme for Lac and MMA. Inversion profiles of the editing pulses applied in the four sub-experiments A-D. HERMES acquires all four combinations of (ON_Lac_, OFF_Lac_) and (ON_MMA_, OFF_MMA_).

The editing scheme and the inversion profiles of the editing pulses are shown in Figure 1b. Sub-experiment A employs a cosine-sinc-Gaussian inversion pulse for dualband inversion, while sub-experiments B and C use a single-lobe sinc-Gaussian pulse. In sub-experiment D (OFF_Lac_, OFF_MMA_), which does not require an editing pulse, a single-lobe pulse is applied at 7.22 ppm.

Since the encoding relies on mutually orthogonal columns of a Hadamard matrix and the separation of the editing targets is larger than the editing pulse bandwidth, the MMA-edited spectrum should contain no edited signal from Lac, and vice versa.

### 2.2 Density-matrix simulations

Density-matrix simulations of the MMA and Lac spin systems were carried out in FID-A^11^, as implemented in MRSCloud^12^. Simulated sub-experiments were combined using Hadamard encoding.

### 2.3 MR experiments

Phantom and *in vivo* data were acquired on a Philips 3T scanner, using a 32-channel head coil. The bandwidths of the slice-selective excitation and refocusing pulses were 2.2 kHz and 1.3 kHz, respectively. The editing pulse duration and bandwidth (defined as the full width at half maximum of each editing lobe) were 30 ms and 41.3 Hz, respectively. Additional acquisition parameters are included in the Minimum Reporting Standards in MRS^13^ table, included as Supplementary material.

#### 2.3.1 Phantom HERMES

Three phantoms were prepared in phosphate-buffered saline (with 1.5 g/l NaN_3_) with: 5 mM MMA; 5 mM Lac; and 5 mM MMA + 5 mM Lac. HERMES experiments were performed on each phantom. Acquisition parameters included: 3 × 3 × 3 cm^3^ voxel; TE/TR 140/2000 ms; 30-ms editing pulses; 16 total transients, and Philips ‘excitation’ water suppression.

#### 2.3.2 *In vivo* HERMES of MMA and Lac

##### Participants

one healthy infant (male, age 1.1 years) and one patient with MMAemia (male, age 10 years) were recruited with approval of the local Institutional Review Board and informed parental consent and subject assent to participate. For demonstration of the method in a healthy subject, an infant was chosen because MMA levels are expected to be negligible, while Lac levels, although low, may be slightly higher than in older healthy subjects^14,15^.

##### MRS

HERMES data were acquired from three voxel regions (36 × 36 × 36 mm^3^) in: basal ganglia^16^; left frontal lobe; and left parietal lobe^17^. All *in vivo* experiments were performed as for the phantom data, except that 160 total transients were acquired (i.e., 40 transients each for sub-experiments A, B, C and D), and VAPOR water suppression^18^ was used. Four water reference transients were acquired without editing pulses or water suppression for interleaved water referencing (IWR), allowing F_0_ to be updated to maintain stable editing^19^. The scan duration for each measurement was 5 minutes 30 seconds.

### 2.5 Post-processing

All data were processed using the open-source MRS data processing and analysis toolbox Osprey 2.9.0^20^ within MATLAB R2024a. The analysis procedures followed consensus-recommended processing guidelines for MRS data^21,22^. Briefly, standardized MRS processing steps in Osprey include eddy-current correction based on unsuppressed water reference^23^, frequency- and phase-correction using probabilistic spectral alignment^24^, removal of residual water signals using a Hankel singular value decomposition (HSVD) filter^25^, and linear-combination modeling of the metabolite spectra. A nine-metabolite custom basis set was generated using MRSCloud with simulated profiles for creatine Cr, glycerophosphocholine GPC, glutamine Gln, glutamate Glu, *myo*-inositol mI, Lac, MMA, N-acetyl aspartate NAA, and phosphocholine PCh. The modeled spectral range was from 0.2 to 4.2 ppm, and a relatively rigid baseline knot spacing of 0.55 ppm^26^ was used. Water-reference data were modeled in the frequency domain with a dedicated water basis function. The levels of MMA and Lac were estimated from their respective difference-edited spectra. Metabolite amplitude estimates were reported as ratios to unsuppressed water in each voxel.

## 3. Results

### 3.1 HERMES density-matrix simulations

Density-matrix simulations of MMA and Lac are shown in Figure 2. The MMA signal at 1.23 ppm is refocused in sub-experiments A and B (ON_MMA_) and inverted in sub-experiments C and D (OFF_MMA_), due to free evolution of the *J*-coupling. Correspondingly, the Lac signal at 1.33 ppm shows a refocused doublet in sub-experiments A and C (ON_Lac_) and an inverted doublet in sub-experiments B and D (OFF_Lac_). The two Hadamard combinations of these sub-experiments, shown in Figure 2, have edited signals in the intended Hadamard-combination spectra, and no crosstalk to the other spectrum, i.e., the MMA simulation gives signal in the MMA-edited difference spectrum and not in the Lac-edited difference spectrum.

**Figure 2.**
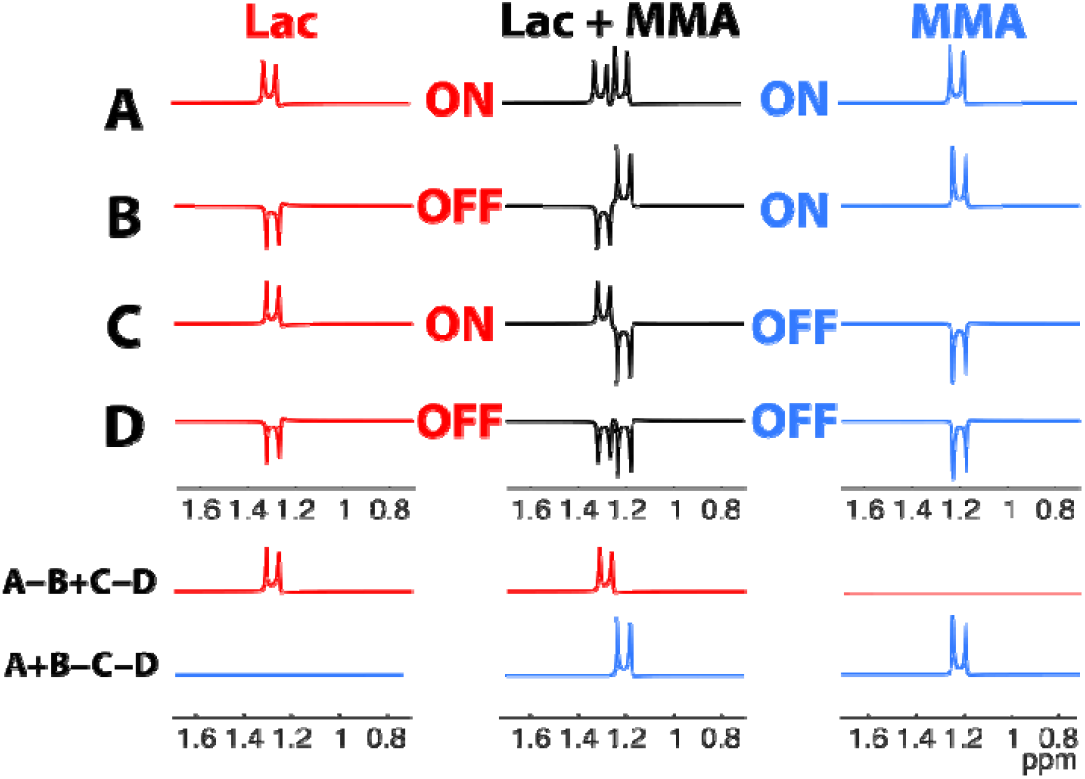
HERMES simulations. Rows A, B, C, and D represent sub-experiments: A ON_Lac_, ON_MMA_; B OFF_Lac_, ON_MMA_; C ON_Lac_, OFF_MMA_; and D OFF_Lac_, OFF_MMA_. MMA and Lac have structural similarities, resulting in a Lac peak is at 1.33 ppm (in red) and an MMA peak at 1.23 ppm (in blue).

### 3.2 Phantom HERMES data

The four HERMES subspectra from the three phantoms (containing MMA-only, Lac-only, and MMA+Lac), are shown in Figure 3. Sub-experiments A-D show the expected doublet patterns, matching the simulations (with slightly wider experimental linewidth). The Hadamard combinations for the three phantoms also strongly resemble the equivalent simulations with crosstalk close to the noise floor. The results of HERMES editing in the phantom that contains both MMA and Lac demonstrate very good separation of the simultaneously acquired signals, again with strong agreement with the simulations.

**Figure 3.**
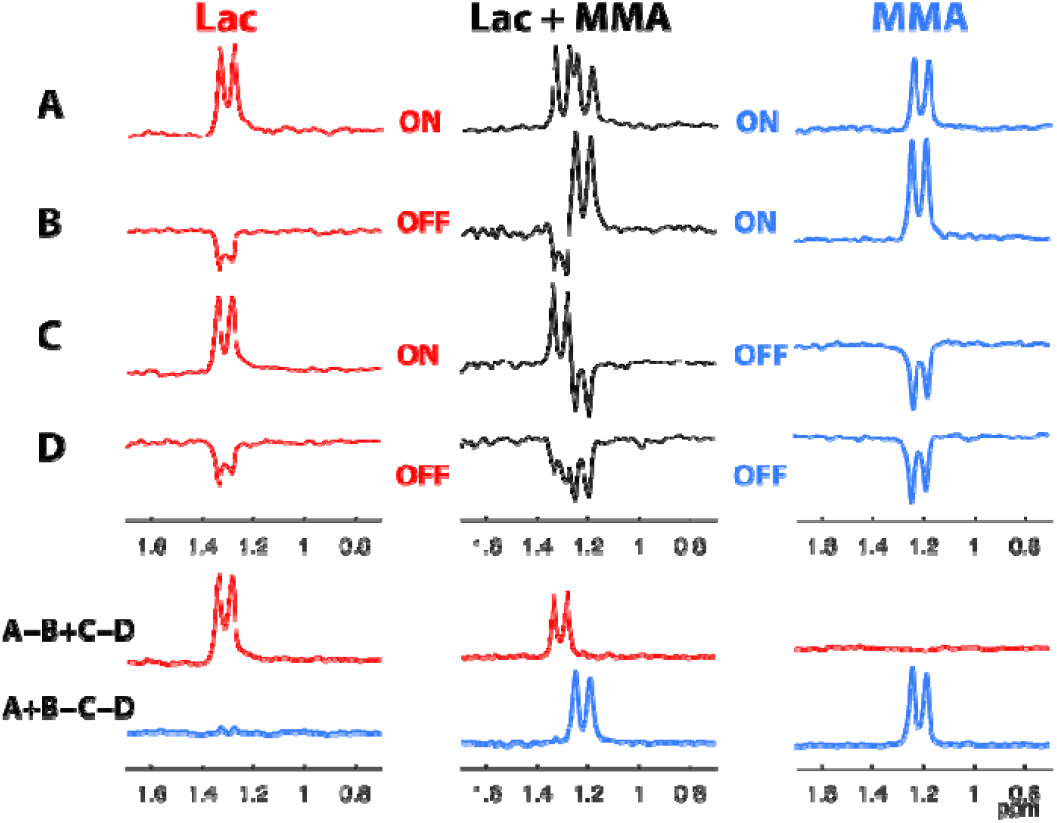
Phantom HERMES experiments. Rows A, B, C, and D represent sub-experiments: A ON_Lac_, ON_MMA_; B OFF_Lac_, ON_MMA_; C ON_Lac_, OFF_MMA_; and D OFF_Lac_, OFF_MMA_. The sum of the two ON_Lac_ sub-experiments minus the sum of the two OFF_Lac_ sub-experiments (A-B+C-D) gives the Lac-edited difference spectrum (red row). The MMA-edited difference spectrum is calculated by subtracting the two OFF_MMA_ sub-experiments from the sum of the two ON_MMA_ sub-experiments (A+B-C-D) (blue row).The separation of signals from the mixed MMA + Lac phantom into distinct Hadamard difference spectra is excellent.

### 3.3 *In vivo* HERMES data

Data from one healthy infant (frontal lobe voxel) is shown in Figure 4, and one MMAemia patient (3 voxels) is shown in Figure 5. The four HERMES subspectra are shown separately in Figure 4a to demonstrate that the editing pulses operated as intended (the choline signal being suppressed in ON_MMA_ subspectra and the creatine signal at 3.9 ppm impacted in ON_Lac_ subspectra). The Hadamard-combination spectra, shown in Figure 4b, show a small Lac signal in the Lac-edited spectrum and no MMA signal in the MMA-edited spectrum (of the healthy subject). The estimated concentration of Lac was 1.42, 1.20 and 1.63 i.u. in the basal ganglia, frontal lobe and parietal lobe, respectively. Linear combination models did not detect an MMA component in any of the three voxels in the healthy infant, as levels were below the detection threshold. Figure 5 shows the Hadamard-combination edited spectra for all three regions in a patient with MMAemia, showing both visible Lac and MMA signals. The estimated concentration of Lac was 1.37, 1.31, and 1.01 i.u. in the basal ganglia, frontal lobe, and parietal lobe, respectively. The estimated concentration of MMA was 0.95, 1.58, and 0.94 i.u. in the basal ganglia, frontal lobe, and parietal lobe, respectively.

**Figure 4.**
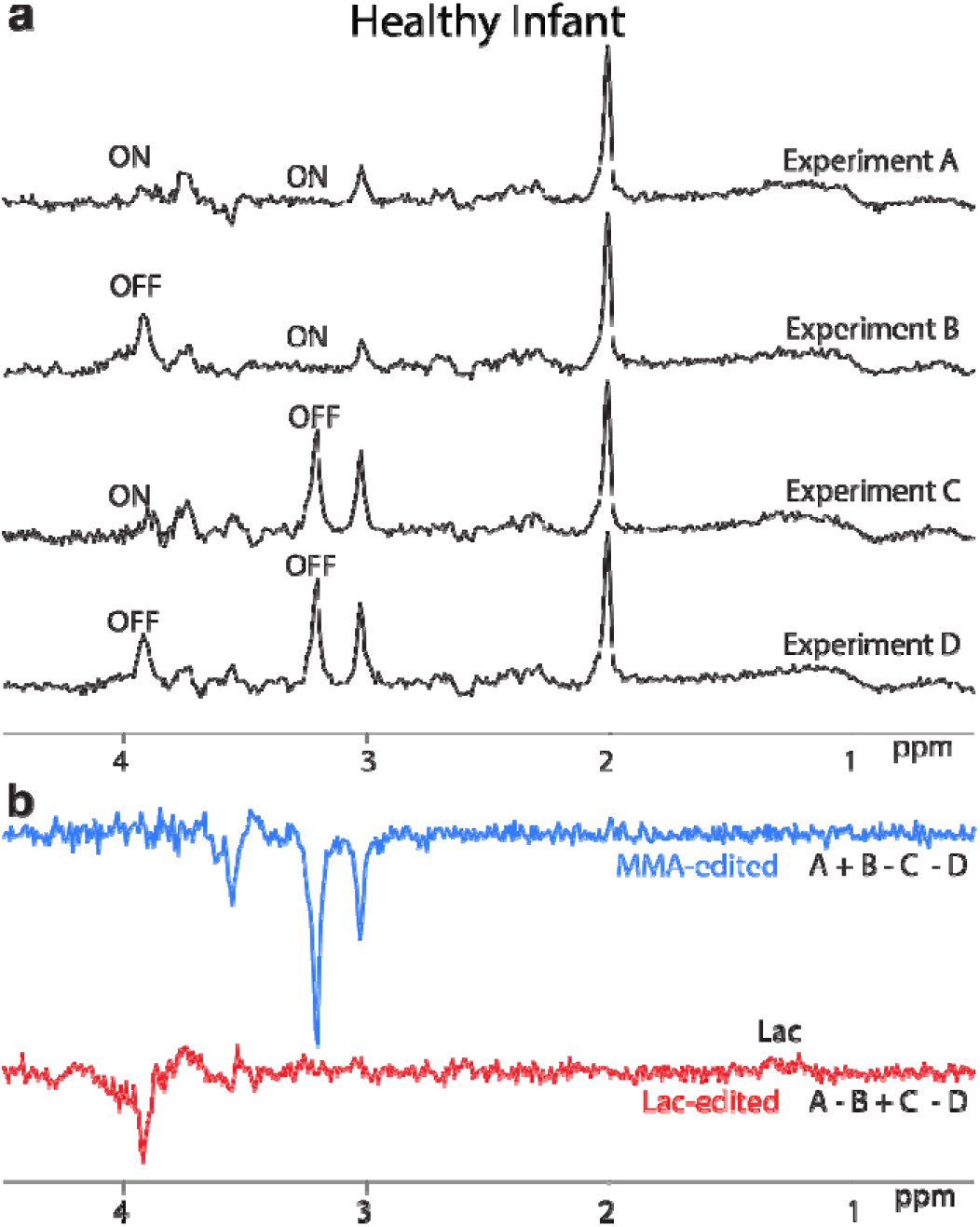
*In vivo* HERMES experiments from a frontal lobe voxel (36 × 36 × 36 mm^3^) of one healthy infant (M, 1.1 years old). a. Separate plots of the HERMES sub-spectra A-D. b. The Hadamard-combination difference spectra yield an MMA-edited spectrum (without MMA signal, expected for a healthy control) in the A+B-C-D combination (blue panel), and a small Lac-edited signal at 1.33 ppm in the A-B+C-D combination (red panel).

**Figure 5.**
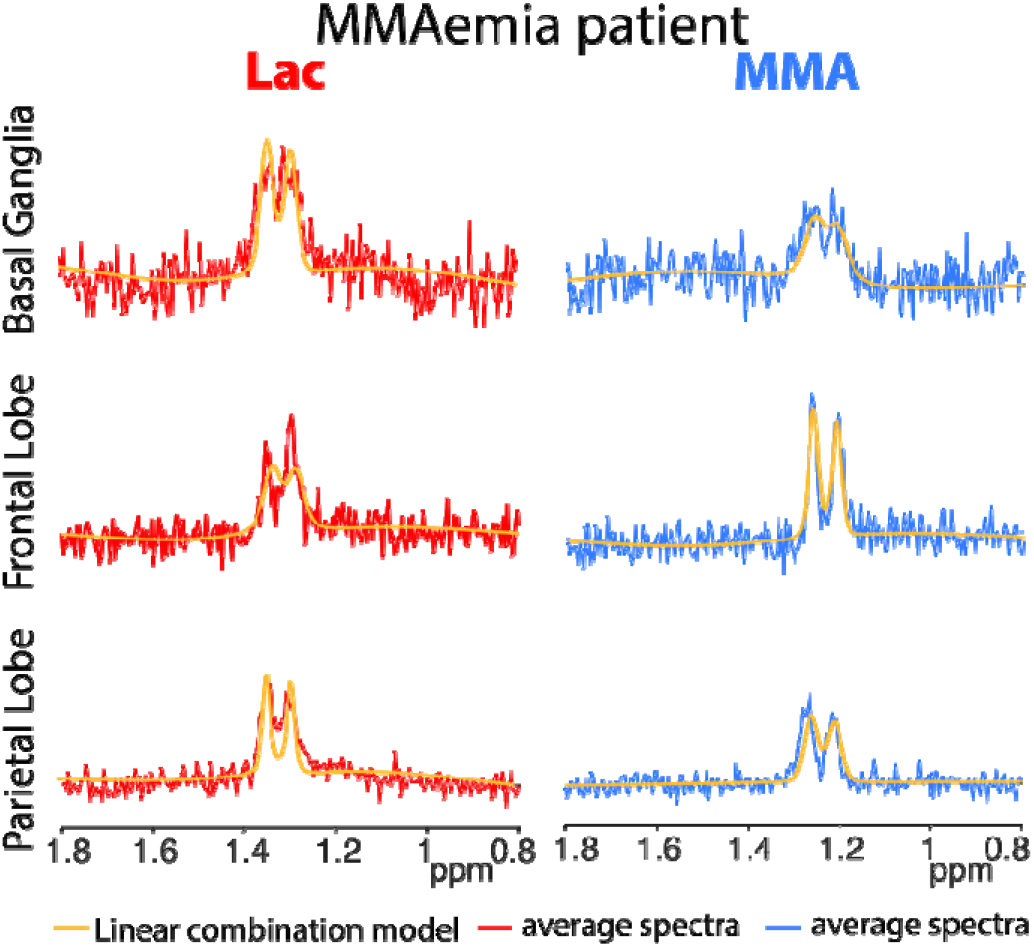
*In vivo* HERMES spectra from one MMAemia patient (M, 10 years old). Lac and MMA peaks are shown in all three regions of interest, resolved in the respective Hadamard-reconstructed difference spectrum. Red/blue lines represent the average spectra, and yellow lines represent the Osprey linear-combination model.

## 4. Discussion

The edited separation of MMA and Lac allows for the distinct and quantitative analysis of two overlapping brain metabolites, in preparation for studies investigating the relationship between brain chemistry and cognitive and clinical symptoms in MMA accumulation disorders. The signals of MMA overlap with other signals in the spectrum (e.g., lipid and Lac), making quantification challenging. The MR spectrum of an individual with MMAemia often displays a increased signals around 1.3 ppm, historically assigned to Lac^10^. Our novel HERMES editing method successfully separates MMA and Lac and demonstrates that both can be present in MMAemia. Simulations and phantom experiments of this HERMES editing protocol demonstrate effective separation of the edited signals into their respective MMA and Lac subspectra, with little crosstalk. *In vivo*, the method was effectively applied in a healthy infant (where no MMA signal and low Lac were observed, as expected) and a patient with MMAemia, in which both Lac and MMA were detected. In practical terms, HERMES enables the separate edited measurement of both MMA and Lac in a single 5-minute acquisition.

The MRS literature of MMAemia is limited. One case report^27^ in a 4-year-old with elevated blood MMA reports low NAA levels, and a signal at 1.3 ppm corresponding to “less than 2.5 mM of lactate … probably coming from lipids”. Trinh et al^10^ report spectroscopic imaging on two cases, one with elevated lactate in basal ganglia lesions, and one (with no lesions) showing elevated Lac in CSF areas. More recently, at 3T^28^, it has been shown that frequency-selective editing can selectively modulate the MMA and Lac signals in patients with MMAemia. In that study, a 20-ms editing pulse was applied at 3.2 ppm to target MMA without impacting the Lac signals. This was implemented as an “edit-on-only” protocol without difference editing, so that MMA and Lac signals appear with opposite polarity. However, some degree of ambiguity remains due to spectral overlap between the two signals, and lipid contamination may complicate quantification further. HERMES uses *J*-difference editing to separate the MMA and Lac signals, and Hadamard encoding to do so simultaneously in a single experiment.

The neurobiological mechanisms by which MMA affects the brain are not well understood. MMAemic mutations disrupt the primary catabolic pathway for branched-chain amino acids^1,29^ (and other molecules such as odd-chain fatty acids), leading to the accumulation of methylmalonyl-CoA. This yields elevated MMA and propionyl CoA, and downstream increases in propionic acid (PA) and 2-methylcitric acid (2-MCA)^2^. These accumulating intermediates inhibit multiple enzymes of the TCA cycle^2^, shifting metabolism toward glycolysis. MMA also inhibits glutamine synthase, leading to elevated ammonia levels^2^ that further suppress several TCA-cycle enzymes and inhibit cytosolic lactate dehydrogenase. This increases Lac levels and impairs neuronal use of Lac as a TCA cycle substrate^29^. Thus, collectively, products elevated as a result of MMAemic mutations reduce TCA cycle turnover, a critical engine of cellular metabolism in terms of energy production and metabolic intermediates. Patients with MMAemia show elevated MMA concentration systemically^30,31^. This mechanistic cascade provides a biochemical model of MMAemia explaining the increased MMA and Lac levels observed systemically, and in the brain, and outlining how they impact cellular function and brain development.

There are some limitations in this study. Its scope as a methodological proof-of-principle study is limited to demonstrating the technical efficacy of HERMES editing for MMA and Lac, and therefore we have only presented data from one patient and one healthy infant. These demonstrate that the method works for editing Lac with or without MMA present, and for editing MMA if it occurs or low signal when it does not. The scope of this study does not include a statistically meaningful cohort from which to draw scientific conclusions, either about the frequency of occurrence of MMA signal in the brain in MMAemia, or comparisons of the level of Lac between patients and controls. Such a study is planned for the future, once the methodology has undergone rigorous peer review. A second limitation is the compromise taken on scan duration - 160 transients were acquired in a 5½-minute scan, which may be insufficient to produce high-quality edited spectra for low-concentration targets. This decision was taken to improve compliance (without anesthesia) during an experiment designed for pediatric research in a group that typically have intellectual disabilities. Practically, this means that there is some detection floor below which MMA (and Lac) may be present but not detected. However, since this experiment will mostly be used for monitoring progression and treatment in MMAemia, rather than for detection and diagnosis, this represents an acceptable compromise. Thirdly, *J*-difference editing as an approach has some inherent limitations. Subtracting two large signals to reveal a small one requires stability – either of the experiment and subject, or of the data after post-processing. Hadamard-edited scans like HERMES require the addition/subtraction of four interleaved sub-experiments instead of two and are therefore even more demanding. Thus, subject motion, incompletely mitigated by post-processing, can result in subtraction artefacts that can bias metabolite quantification^32^. Beyond subtraction artefacts, the use of relatively narrow frequency-selective editing pulses requires stable B_0_ for efficient editing. Both scanner drift and subject motion can cause instability in B_0_, although this is mitigated by real-time frequency F_0_ updates by IWR. Despite this limitations, HERMES-edited MRS has been applied in pediatric contexts previously, including in studies of developmental disorders^33–35^. A final limitation is the finite selectivity of editing and the potential for signal overlap. Using an editing pulse centered at 4.1 ppm will co-edit threonine^36^ (4.25 ppm) and β-hydroxybutyrate^37,38^ (bHB, 4.13 ppm) where present. Their edited methyl signals at 1.32 ppm and 1.19 ppm respectively may complicate Lac quantification.

Additionally, the Lac doublet at 1.33 ppm sits on a broad background of macromolecular and lipid methyl resonances ∼1.2-1.4 ppm, which can co-edit^39,40^, but also generate substantial subtraction artefacts. Including additional components in the basis set in future studies may mitigate signal overlap.

## 5. Conclusion

In conclusion, we have presented a new application of HERMES to separate the overlapping signals of MMA and Lac. This experiment is likely to be useful for studies investigating the relationship between MMA levels and intellectual disability, between MMA levels centrally and systemically, and changes in MMA with diet modification and other potential treatments.

## Acknowledgements

This work was supported by National Institutes of Health (NIH) grants R01 EB032788, R01 EB016089, R01 EB023963, R01 EB035529, R21 EB033516, P41 EB031771, and K99 HD118185. This work was supported by the National Natural Science Foundation of China [No. 82302149 (TG); U25A20137(TG)], Shandong Technological Innovation Guidance Program [No. YDZX2025033(GW)], Shandong Provincial Natural Science Foundation [No. ZR2020QH267(TG); ZR2024LSW010(TG); ZR202211210106 (GW)],China Postdoctoral Science Foundation [No. 2022M711987 (TG)].

