## Supplementary material for "Methylmalonic acid: a new target for Hadamard-edited MRS"

| 1.Hardware | |
| --- | --- |
| a. Field strength [T] | 3 T |
| b. Manufacturer | Philips |
| c. Model (software version if available) | R 5.7.1 |
| d. RF coils: nuclei (transmit/receive), number of channels, type, body part | ^1^H, 32 channel, head |
| e. Additional hardware | - |
| 2. Acquisition | |
| a. Pulse sequence | HERMES (Johns Hopkins University Patch) |
| b. Volume of interest (VOI) locations | Basal ganglia, frontal lobe, parietal lobe |
| c. Nominal VOI size [mm^3^] | 36 x 36 x 36 mm^3^ |
| d. Repetition time (TR), echo time (TE)[ms] | TR 2000 ms, TE 140 ms |
| e. Total number of averages per spectrum  i. Number of averaged specra per subspectrum | 160 total averages with 40 averages per subspectrum |
| f. Additional sequence parameters  i. Editing pulse parameters | F1: 2000 Hz, 2048 points  MMA at 3.2 ppm, Lac at 4.1 ppm |
| g. Water suppression method | VAPOR |
| h. Shimming method, reference peak, and threshold of acceptance of shim chosen | 1^st^ and 2^nd^ shimming pencil beam, water |
| i. Trigger or motion correction | No trigger or active motion correction |
| 3. Data analysis methods and outputs | |
| a. Analysis software | Osprey 2.9.0 |
| b. Processing steps deviating from Osprey | Final alignment of the averaged sub-spectra by  minimizing the choline (not water) peak; |
| c. Output measure | tCr, rawWaterScaled |
| d. Quantification references and assumptions, fitting model assumptions | NAA, Cr, GPC, PCh, Lac, MMA, mI, Glu, Gln Fitting method: Osprey baseline knot spacing 0.55 ppm |
| 4. Data quality | |
| HERMES, parietal lobe | |
| a. SNR (Cr), linewidth (Cr) [Hz, OFF spectra] | SNR 55.93, linewidth 4.00 Hz |
| c. Quality measures of postporcessing model fitting (Mean Relative Amplitude Residual (Residual/Noise))  sum  diff1  diff2 | 5.07  3.03  9.39 |
| HERMES, frontal lobe | |
| a. SNR (Cr), linewidth (Cr) [Hz, OFF spectra] | SNR 50.98, linewidth 4.46 Hz |
| c. Quality measures of postporcessing model fitting (Mean Relative Amplitude Residual (Residual/Noise))  sum  diff1  diff2 | 7.50  1.56  1.81 |
| HERMES, basal ganglia | |
| a. SNR (Cr), linewidth (Cr) [Hz, OFF spectra] | SNR 67.61, linewidth 3.48 Hz |
| c. Quality measures of postporcessing model fitting (Mean Relative Amplitude Residual (Residual/Noise))  sum  diff1  diff2 | 5.51  1.55  1.42 |

Supplementary Table 1: Summary following minimum reporting standards in MRS generated in Osprey See Lin et al. 'Minimum Reporting Standards for in vivo Magnetic Resonance Spectroscopy (MRSinMRS): Experts' consensus recommendations. NMR in Biomedicine. 2021;e4484. [doi.org/10.1002/nbm.4448](https://doi.org/10.1002/nbm.4448)
